# *Toxoplasma gondii* importin α shows weak auto-inhibition

**DOI:** 10.1101/2022.10.06.510747

**Authors:** Manasi Bhambid, Vishakha Dey, Sujata Walunj, Swati Patankar

## Abstract

Importin α is a nuclear transporter that binds to nuclear localization signals (NLSs), consisting of 7-20 positively charged amino acids found within cargo proteins. In addition to cargo binding, intramolecular interactions also occur within the importin α protein due to binding between the importin β-binding (IBB) domain and the NLS-binding sites, a phenomenon called auto-inhibition. The interactions causing auto-inhibition are driven by a stretch of basic residues, similar to an NLS, in the IBB domain. Consistent with this, importin α proteins that do not have some of these basic residues lack auto-inhibition; a naturally occurring example of such a protein is found in the apicomplexan parasite *Plasmodium falciparum*. In this report, we show that importin α from another apicomplexan parasite, *Toxoplasma gondii*, harbors basic residues (KKR) in the IBB domain and exhibits auto-inhibition. This protein has a long, unstructured hinge motif (between the IBB domain and the NLS-binding sites) that does not contribute to auto-inhibition. However, the IBB domain may have a higher propensity to form an α-helical structure, positioning the wild-type KKR motif in an orientation that results in weaker interactions with the NLS-binding site than a KRR mutant. We conclude that the importin α protein from *T. gondii* shows auto-inhibition, exhibiting a different phenotype from that of *P. falciparum* importin α. However, our data indicate that *T. gondii* importin α may have a low strength of auto-inhibition. We hypothesize that low levels of auto-inhibition may confer an advantage to these important human pathogens.

## INTRODUCTION

Eukaryotic cells are defined by nuclear and cytoplasmic compartments separated by the nuclear envelope. Due to this compartmentalization, the transport of nuclear proteins occurs through nuclear pore complexes (NPCs) that perforate the nuclear envelope (Bonner, 1975; Feldherr et al., 1983). Nuclear proteins bear nuclear localization signals (NLSs) and are actively transported by carrier proteins, importin α and importin β (Adam et al., 1990; Cassany & Gerace, 2009; Dingwall & Laskey, 1991). Importin α recognizes classical NLSs, characterized by clusters of basic amino acids, while importin β facilitates the passage of the importin α:cargo complex by interactions with the NPC (Conti et al., 1998; Görlich et al., 1995).

The importin α structure consists of NLS-binding sites within a tandem series of Armadillo (ARM) repeats and an N-terminal importin β-binding (IBB) domain (Görlich et al., 1996; Conti et al., 1998). The structurally polymorphic IBB domain mimics positively charged NLS sequences (Fanara et al., 2000; Jibiki et al., 2021; Lott & Cingolani, 2011) and binds to the ARM repeats of importin α, thereby blocking the NLS-binding sites, a phenomenon called auto-inhibition. Auto-inhibition has been mapped to the third of the three basic amino acid clusters in the IBB domain (Harreman et al., 2003; Kobe, 1999). The IBB domain binds to importin β, revealing the NLS-binding sites and relieving auto-inhibition. Between the IBB domain and the ARM repeats lies a hinge motif that provides flexibility for these auto-inhibitory interactions (Kobe, 1999; Sankhala et al., 2017).

Importin α proteins have been studied extensively (Chang et al., 2012; Dey & Patankar, 2018; Harreman et al., 2003; Hu et al., 2005; Hu & Jans, 1999; Hübner et al., 1999; Kobe, 1999; Miyatake et al., 2015; Pumroy et al., 2015; Pumroy & Cingolani, 2015); however, the extent of auto-inhibition of different importin α proteins has not yet been comprehensively compared. Interestingly, full-length *Plasmodium falciparum* importin α (PfImpα) lacks auto-inhibition (Dey & Patankar, 2018), while the full-length importin α of *Toxoplasma gondii* (TgImpα), an apicomplexan parasite like *P. falciparum*, was shown to bind to the NLS of the *T. gondii* histone acetyltransferase, general control non-derepressible 5a (GCN5a), qualitatively suggesting a lack of auto-inhibition (Bhatti & Sullivan, 2005).

In this report, we quantitatively assess and demonstrate that the *T. gondii* importin α protein exhibits auto-inhibition. We systematically test *in vitro* the NLS-binding affinity of mutants in the third basic cluster in TgImpα. Using *in silico* approaches, we predict the structure of native and mutant TgImpα proteins and propose that the helicity of the TgImpα IBB domain may result in weak interactions with the NLS-binding sites. Furthermore, we observe a longer hinge motif in importin α of *T. gondii* and other apicomplexans and test its role in auto-inhibition. This report establishes that, unlike *P. falciparum*, *T. gondii* has an importin α protein that shows auto-inhibition; however, our data indicate that the strength of auto-inhibition may be weak. Hence, importin α proteins from two apicomplexan parasites appear to have evolved towards low levels of auto-inhibition, and we suggest that this phenotype may have a role in the biology of these human pathogens.

## EXPERIMENTAL PROCEDURES

### Cloning of expression constructs

#### Wild-type TgImpα and TgImpα lacking the IBB domain in pET28a

TgImpα (ToxoDB ID: TGGT1_252290) was cloned into the pET28a vector using the NcoI and HindIII sites to generate a fusion with the C-terminal His-tag. All primers in this study were obtained from Sigma-Aldrich, and the restriction sites are underlined. First, cDNA was synthesized using the gene-specific reverse primer. Next, the same reverse primer was used with the forward primer to amplify the 1.64 kb TgImpα gene. The first 92 amino acids from the N-terminus of the TgImpα gene were found to be the IBB domain and hinge of the protein by Pfam analysis (http://pfam.xfam.org/) (Mistry et al., 2021). For generating ΔIBB-TgImpα fused with a His-tag at the C-terminus, DNA corresponding to amino acids 92-545 of the gene was sub-cloned from this plasmid into pET28a between the NcoI and HindIII sites using the respective forward primer and the same reverse primer. The inserts were confirmed to be free of mutations by Sanger sequencing.

Wild-type (forward primer): 5ʹ CATG<u>CCATGG</u>AGCGCAAGTTGGCCGATC 3ʹ

ΔIBB (forward primer): 5ʹ AATA<u>CCATGG</u>GCCTCTCCAGCGGAGATCCG 3ʹ

Reverse primer: 5ʹ TCCC<u>AAGCTT</u>CTGGCCGAAGTTGAAGCCTC 3ʹ

#### TgImpα third basic residue cluster mutants

The KKR motif was mutated into SKR, KRR and AAA by site-directed mutagenesis. The PCR products were amplified with a forward primer designed to carry the desired mutation (underlined) and a common reverse primer using WT-TgImpα with the C-terminal His-tag template. The mutations were confirmed by Sanger sequencing.

KKR/SKR (Forward primer): 5ʹ CAGAACTTGGCA<u>TCGAAGCGC</u>GCGGAGGCGCTG 3ʹ

KKR/KRR (Forward primer): 5ʹ CAGAACTTGGCA<u>AAGCGCCGC</u>GCGGAGGCGCTG 3ʹ

KKR/AAA (Forward primer): 5ʹ CAGAACTTGGCA<u>GCTGCTGCT</u>GCGGAGGCGCTG 3ʹ

Reverse primer: 5ʹ CTCGCGGTGCGTCTTGCGAATCTGCAGCTGCAG 3ʹ

#### TgImpα chimera containing the hinge motif from MmImpα2

The hinge motif in TgImpα (amino acids 49-78) was substituted with amino acids 57-70 of the hinge motif from MmImpα2 (Uniprot ID: P52293). This construct (MmHinge-TgImpα) was generated using primers with a sequence coding for the MmImpα2 hinge (underlined) and sequence overlap with the TgImpα gene. WT-TgImpα in the pET28a vector with the C-terminal His-tag template was used. The sequence of the chimera was confirmed by Sanger sequencing.

MmHinge (Forward primer): 5ʹ <u>CTACAGGAAAACCGGAACAAC</u>AACGTCTTCAGCTTCGAACATC 3ʹ

MmHinge (Reverse primer): 5ʹ <u>CGGAGAAGTAGCATCATCAGG</u>GTCCAGCGCCTCCGC 3ʹ

### Binding assays to study importin α:NLS interaction

#### Gel-based binding with Ni-NTA beads

Freshly purified WT-TgImpα and ΔIBB-TgImpα proteins were used as bait, while the NLS from the *P. falciparum* trimethyl guanosine synthase 1 (PfTGS1) protein was used as prey. Thrombin treatment was used to remove the His-tag from the His-tag fused PfTGS1-NLS-GFP and GFP as described previously (Babar et al., 2016). Equimolar concentrations of WT-TgImpα and ΔIBB-TgImpα (bait) proteins (2.5 μM) were incubated separately with His-tag-free PfTGS1-NLS-GFP and GFP (prey) proteins (5 μM). The reactions were carried out using bait bound to Ni-NTA beads as described previously (Dey & Patankar, 2018) to pull down the prey protein. The beads were then washed and added to SDS-PAGE gel loading dye to prepare samples before running on 15% SDS-PAGE gel. The gels were stained with Coomassie Blue to visualize the bands using standard protocols. The experiment was repeated twice. The intensity of the PfTGS1-NLS-GFP bands was quantified using the Gel Analysis method in ImageJ software (Schneider et al., 2012; Stael et al., 2022). The relative intensity is a ratio of the % intensity of the PfTGS1-NLS-GFP band pulled down by ΔIBB-TgImpα compared to that pulled down by WT-TgImpα.

#### Surface Plasmon Resonance (SPR)

The SPR assay was performed using the T200 BIAcore instrument described previously (Dey & Patankar, 2018). Due to the stability of the analytes and ligands, they were either freshly prepared or stored at −80°C. Analytes (WT-TgImpα and mutants) were prepared at the concentration range by a series of 2-fold dilutions and were allowed to interact with the immobilized ligands (PfTGS1-NLS-GFP and GFP) on a CM5 sensor chip. The binding was calculated after blank subtraction from ligand-coated chips interacting with GFP. The Kd was determined by fitting the sensorgrams in a global 1:1 Langmuirien interaction model, using BIA evaluation software 1.0 (GE Healthcare) and plotted in GraphPad Prism 9.0.2. All kinetic assays were performed three times.

#### AlphaScreen

The AlphaScreen binding assay was performed as described previously (Fraser et al., 2014; Thomas et al., 2018; Wagstaff et al., 2011) to study interactions between TgImpα (with or without *M. musculus* importin β1) with SV40 T-ag-NLS-GFP (Walunj et al., 2022). The previously published plasmid expressing WT-TgImpα with a C-terminal Transferase (GST) tag was used for the assay (Walunj et al., 2022). The SV40 T-ag-NLS-GFP and MmImpβ1 plasmids were the same that those published previously (Wagstaff et al., 2012; Wagstaff & Jans, 2006). GST-tagged TgImpα and MmImpβ1 were pre-dimerized as previously described (Wagstaff et al., 2012). For the binding assay (performed at room temperature), 30 nM final concentration of SV40-NLS-GFP-His was added to each well, followed by the addition of TgImpα or pre-dimerized TgImpα/MmImpβ1 (0-15 nM final concentration range) and incubation for 30 minutes. Then, 4 μL of the acceptor beads (Perkin Elmer) diluted in PBS was added to each well in the dark and incubated for 90 mins. Next, 4 μL of the donor beads (Perkin Elmer) diluted in PBS was added to each well to give a total volume of 25 μL and incubated for 2 hours. The AlphaScreen signal was quantified on a Perkin Elmer plate reader, and titration curves (sigmoidal fit) were plotted using GraphPad Prism 9.0.2 (San Diego, California, USA) as previously mentioned (Walunj et al., 2022). The hooking zone values were excluded, reflecting the signal’s quenching due to the oversaturation of one of the beads’ binding partners (Wagstaff et al., 2012).

### Structure prediction and docking

*Ab initio* structures of the full-length TgImpα (Uniprot ID: A0A125YID0), the TgIBB domain (amino acids 1-49) and TgARM repeats with the hinge (amino acids 50-545) and the full-length MmImpα2 (Uniprot ID: P52293) were generated using the RaptorX Contact Prediction Server (http://raptorx.uchicago.edu/) (Källberg et al., 2012). The predicted models were corroborated in the Robetta structure prediction program (https://robetta.bakerlab.org/) and the AlphaFold protein structure database (https://alphafold.ebi.ac.uk/) (Baek et al., 2021; Jumper et al., 2021; Varadi et al., 2022). The stereochemical quality of the predicted models from RaptorX (with the best RMSD value) was checked on the PROCHECK server (https://saves.mbi.ucla.edu/) (R. A. Laskowski et al., 1993; RomanA. Laskowski et al., 1996). The helical content of the predicted structures was determined in the Agadir program (http://agadir.crg.es/protected/academic/calculation.jsp) (Lacroix et al., 1998; Muñoz & Serrano, 1994, 1995a, 1995b, 1997).

The predicted IBB domain structure and ARM repeat structure were docked in the HADDOCK 2.4 docking server (https://wenmr.science.uu.nl/haddock2.4/) (Honorato et al., 2021; van Zundert et al., 2016; Wassenaar et al., 2012). The residues directly involved in the interaction (as observed in MmImpα2; PDB ID: 1IAL) (Kobe, 1999) were selected for docking in models of TgImpα in the program. The cluster with the lowest Haddock score and RMSD from the overall lowest-energy structure was studied (van Zundert et al., 2016).

### Evolutionary analyses

The evolutionary relatedness between importin α proteins was studied by analyzing the homologs of TgImpα in the plants, animals, fungi, algae, dinoflagellates and alveolates groups. Importin α sequences were selected from the Uniprot database based on nomenclature and the Pfam database based on their domains (ARM, ARM3 and IBB) similar to TgImpα. The dataset includes all importin α proteins from Alveolata groups (Apicomplexa, Perkinsozoa, dinoflagellate, Colpodellida and Ciliata) and some representative genera from the alga, plant, animal and fungi groups. The amino acid sequence of full-length importin α from these organisms was then aligned using the Multiple Sequence Comparison by Log-expectation (MUSCLE) algorithm (Kumar et al., 2018). Sequences that did not align (all dinoflagellates importin α and some sequences from other groups) were discarded. In the multiple Sequence Alignment (MSA) of the 73 proteins, the number of residues in the hinge motif and the proline-glycine amino acids was counted manually. The graphs were plotted using GraphPad Prism 9.0.2 (San Diego, California, USA).

The evolutionary history was inferred using the Maximum Likelihood method and Le_Gascuel_2008 (LG) + Gamma distribution (G) model (Le & Gascuel, 2008). The bootstrap consensus tree inferred from 500 replicates was taken to represent the evolutionary history of the taxa analyzed (Felsenstein, 1985). The percentage of replicate trees in which the associated taxa clustered together in the bootstrap test (500 replicates) are shown next to the branches (Felsenstein, 1985). Neighbor-Join and BioNJ algorithms were applied automatically to obtain the Initial tree(s) for the heuristic search. The matrix of pairwise distances was estimated using the JTT model and then selecting the topology with a superior log-likelihood value.

## RESULTS

### Wild-type *Toxoplasma gondii* importin α demonstrates auto-inhibition

*P. falciparum* importin α shows a lack of auto-inhibition, conferred by a serine residue in the third basic cluster present in the IBB domain (Dey & Patankar, 2018). *T. gondii* is an apicomplexan parasite with a single gene of importin α similar to *P. falciparum*, and the protein encoded by this gene has a KKR motif in the third basic residue cluster. We asked whether TgImpα would show a lack of auto-inhibition, as previously suggested (Bhatti & Sullivan, 2005). To establish the presence or absence of auto-inhibition, it is essential to compare the NLS-binding affinity *in vitro* of (1) a full-length protein to (2) the same protein that lacks auto-inhibition (by either deleting the IBB domain or by including importin β in the reaction). A comparison of the Kd values of these two conditions gives a fold-change value that serves as a quantitative value for auto-inhibition.

A hexapeptide consisting of amino acids 94-99 of TgGCN5a (TGGT1_254555) was functional as a mono-partite NLS *in vivo* (Bhatti & Sullivan, 2005). We attempted to test the binding of TgImpα to this hexapeptide fused to GFP using Ni-NTA agarose beads and found a very faint band resulting from the binding interactions (data not shown). Additionally, in the AlphaScreen assay, the binding affinity of the TgGCN5a-NLS was only marginally higher than that of the negative control (data not shown). Therefore, a strong, bi-partite NLS found in *P. falciparum* trimethyl guanosine synthase 1 (PfTGS1) was used for further experiments. Although derived from a *P. falciparum* protein, this NLS (PfTGS1-NLS) was chosen as it allowed a direct comparison to the strength of auto-inhibition of PfImpα (Dey & Patankar, 2018). Using this NLS, gel-based binding assays were performed to compare the binding affinities of TgImpα and ΔIBB-TgImpα (Figure 1A).

**Figure 1:**
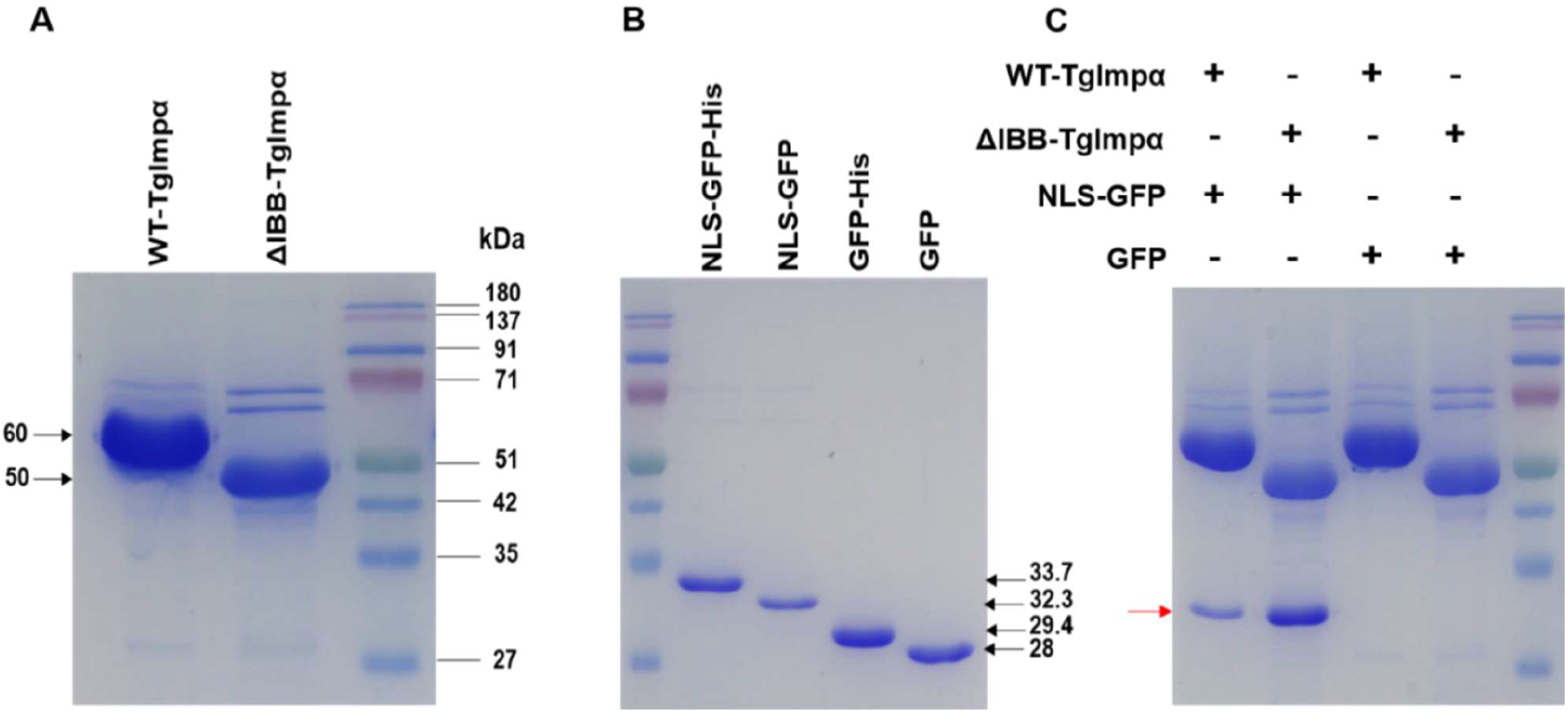
Binding assay using Ni-NTA beads shows that *T. gondii* importin α demonstrates auto-inhibition. **(A)** SDS-PAGE analysis of purified bait proteins for the binding assays. The expected sizes of WT-TgImpα and ΔIBB-TgImpα are 60 kDa and 50 kDa, respectively. **(B)** SDS-PAGE analysis of purified prey proteins. The expected sizes of GFP fused PfTGS1-NLS with the His-tag (PfTGS1-NLS-GFP-His) and without His-tag (PfTGS1-NLS-GFP), GFP with the His-tag (GFP-His) and without His-tag (GFP) are 33.7 kDa, 32.3 kDa, 29.4 kDa and 28 kDa, respectively. **(C)** Gel-based binding assay of the interaction of WT-TgImpα and ΔIBB-TgImpα with PfTGS1-NLS-GFP. The red arrow indicates the prey protein, PfTGS1-NLS-GFP. The assay was done in duplicates (n = 2). Note that WT-TgImpα indicates the full-length TgImpα protein.

Both WT-TgImpα and ΔIBB-TgImpα bound to PfTGS1-NLS-GFP. The intensity of the band corresponding to His-tag free PfTGS1-NLS-GFP (Figure 1B) pulled down by ΔIBB-TgImpα was higher than WT-TgImpα (Figure 1C; indicated by the red arrowhead) by a value of 4.6 ± 1.9 for the replicates. The binding of His-tag-free PfTGS1-NLS-GFP is specific since neither of the bait proteins showed binding to GFP (Figure 1C). These results show that TgImpα exhibits auto-inhibition.

Next, SPR kinetic analysis was performed to identify the difference between the binding affinities of WT-TgImpα (Figure 2A) and ΔIBB-TgImpα (Figure 2B) with PfTGS1-NLS-GFP. A Kd of 0.83 ± 0.04 μM was obtained for WT-TgImpα and 0.24 ± 0.03 μM for ΔIBB-TgImpα, while the negative control (GFP) showed no binding. Consistent with the gel-based binding assays (Figure 1C), quantitative data indicated that ΔIBB-TgImpα exhibits a 3.5-fold higher binding affinity when compared to WT-TgImpα, reconfirming that TgImpα exhibits auto-inhibition.

**Figure 2:**
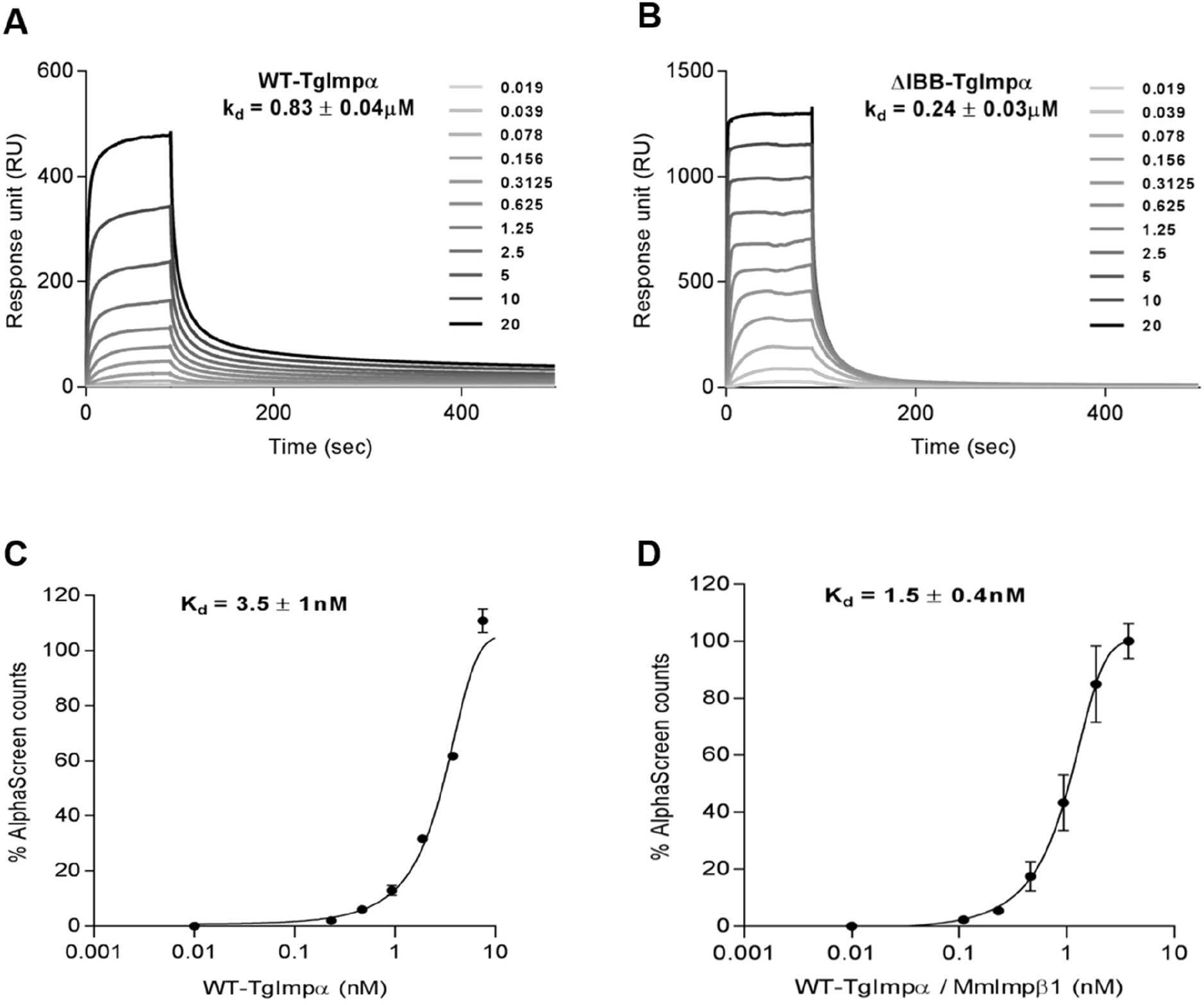
SPR kinetic analysis and AlphaScreen assay confirm that *T. gondii* importin α demonstrates auto-inhibition. SPR kinetic analysis of the interaction of **(A)** WT-TgImpα and **(B)** ΔIBB-TgImpα with PfTGS1-NLS-GFP. The concentration range mentioned is μM. AlphaScreen binding assay of **(C)** WT-TgImpα alone and **(D)** WT-TgImpα/MmImpβ1 dimer with SV40 T-ag-NLS-GFP. All results represent the mean ± SD (n = 3) for Kd values measured. Note that WT-TgImpα indicates the full-length TgImpα protein.

Auto-inhibition was also confirmed using a different approach, the AlphaScreen binding assay (Walunj et al., 2022). Here, a mono-partite NLS (SV40 T-ag-NLS-GFP) was chosen as this NLS has been used extensively (Wagstaff & Jans, 2006). Rather than using ΔIBB-TgImpα, we used the full-length TgImpα protein and included MmImpβ1, which we have shown binds to TgImpα. Perhaps due to the sensitivity of the AlphaScreen technology, a Kd of 3.5 ± 1 nM was obtained for TgImpα alone (Figure 2C) and 1.5 ± 0.4 nM in the presence of MmImpβ1 (Figure 2D). This assay gave a fold-change value of 2.3-fold. Therefore, using three different assays, it can be concluded that TgImpα shows auto-inhibition.

### TgImpα mutants indicate that the third basic residue cluster has a major contribution to auto-inhibition

Multiple Sequence Alignment to compare the IBB domain of TgImpα with that of PfImpα, ScImpα, HsImpα2 MmImpα2, and AtImpα1 showed that the third basic residue cluster of TgImpα has a KKR motif (Figure 3A). To assess whether the third basic residue cluster is solely responsible for the auto-inhibition of TgImpα, we generated three mutants (Figure 3B) where the wild-type KKR motif was replaced with (1) an AAA motif, as seen in the ScImpα mutant lacking auto-inhibition (Harreman et al., 2003), (2) a KRR motif, as seen in importin α proteins that also exhibit auto-inhibition (Harreman et al., 2003; Hu et al., 2005; Hu & Jans, 1999; Hübner et al., 1999; Miyatake et al., 2015) and (3) an SKR motif, as seen in the wild-type PfImpα protein lacking auto-inhibition (Dey & Patankar, 2018).

**Figure 3:**
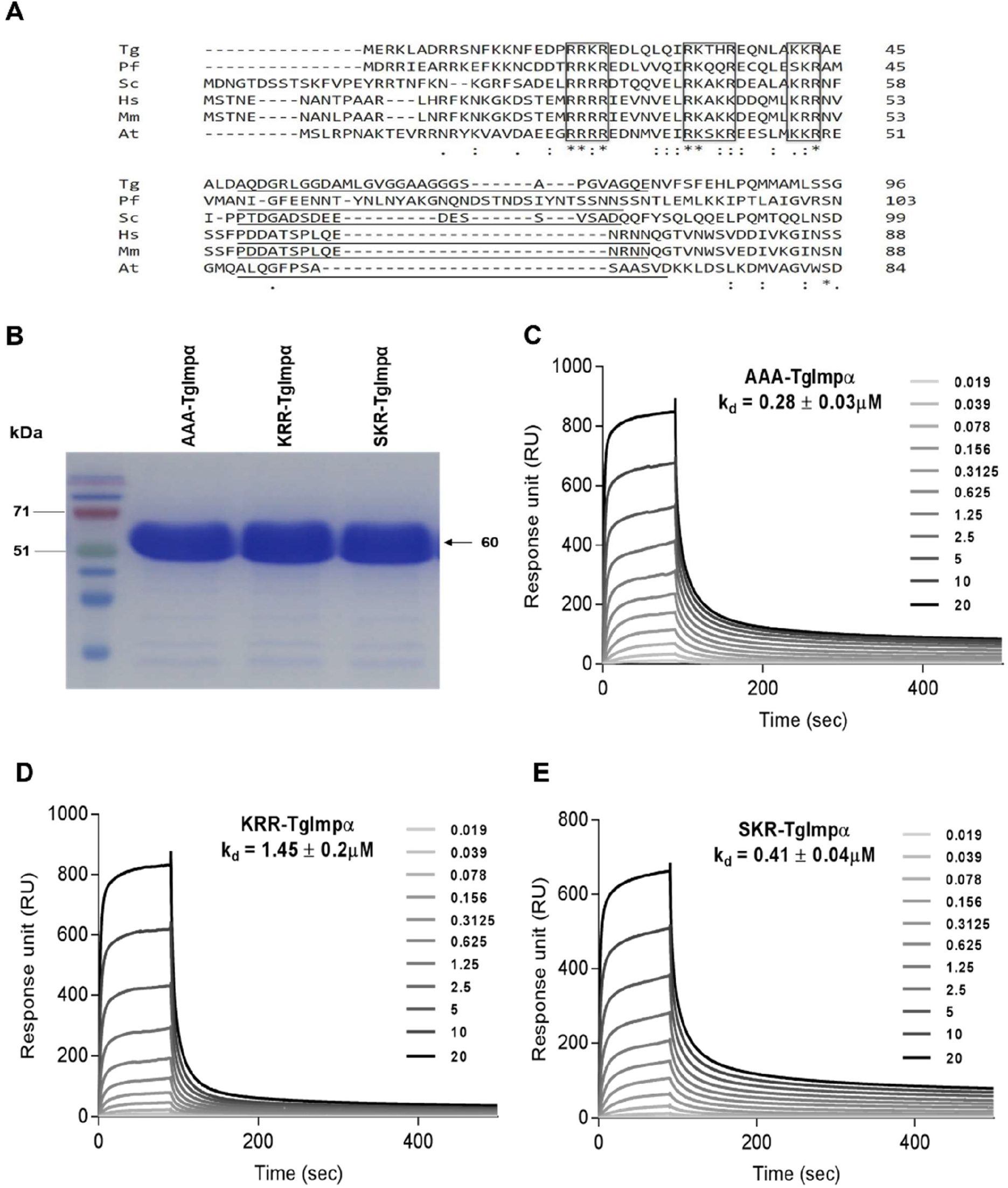
Mutants of TgImpα show differences in auto-inhibition strengths. **(A)** Multiple sequence alignment of importin α proteins using Clustal Omega. Comparison of Tg: *T. gondii* importin α (A0A125YID0) with importin α proteins of Pf: *P. falciparum* (Q8IAW0), Sc: *S. cerevisiae* (A0A6A5Q590), Hs: *H. sapiens* (P52292), Mm: *M. musculus* (P52293) and At: *A. thaliana* (Q96321), with respective Uniprot ID in the bracket. The black boxes represent the basic amino acid clusters in the IBB domain, and the underlined amino acids are the hinge motifs. **(B)** SDS-PAGE of purified TgImpα mutant proteins with the expected sizes of 60 kDa. SPR kinetic analysis of the interaction of **(C)** AAA-TgImpα, **(D)** KRR-TgImpα and **(E)** SKR-TgImpα with PfTGS1-NLS-GFP. All assays were performed in triplicates (n = 3), representing the mean ± SD for Kd values measured. The concentration range mentioned is μM.

Kinetic analysis was performed to determine the Kd of AAA-TgImpα (Figure 3C), KRR-TgImpα (Figure 3D), and SKR-TgImpα (Figure 3E) with PfTGS1-NLS-GFP. All the proteins gave Kd values in the low micromolar range, with AAA-TgImpα showing the highest binding affinity (0.28 ± 0.03 μM), followed by SKR-TgImpα (0.41 ± 0.04 μM) and then KRR-TgImpα (1.45 ± 0.2 μM). These data indicated that mutations in the third basic residue cluster of the TgImpα IBB domain alter the binding strength to the NLS. We next assessed the levels of auto-inhibition by calculating the fold-change of the Kd values of these proteins compared to that of ΔIBB-TgImpα, a protein that reflects no auto-inhibition. We also compared the fold-change values to PfImpα (Figure 4), where systematic mutations of the third basic cluster had been performed (Dey & Patankar, 2018). AAA-TgImpα showed a lack of auto-inhibition, as seen by a 1.2-fold increase in Kd, compared to ΔIBB-TgImpα. Consistent with our previous report, the SKR mutant of TgImpα also showed a lack of auto-inhibition like *P. falciparum* importin α (a 1.7-fold increase compared to ΔIBB-TgImpα) (Dey & Patankar, 2018). These data suggested that the third basic residue cluster plays a major role in the auto-inhibition of TgImpα.

**Figure 4:**
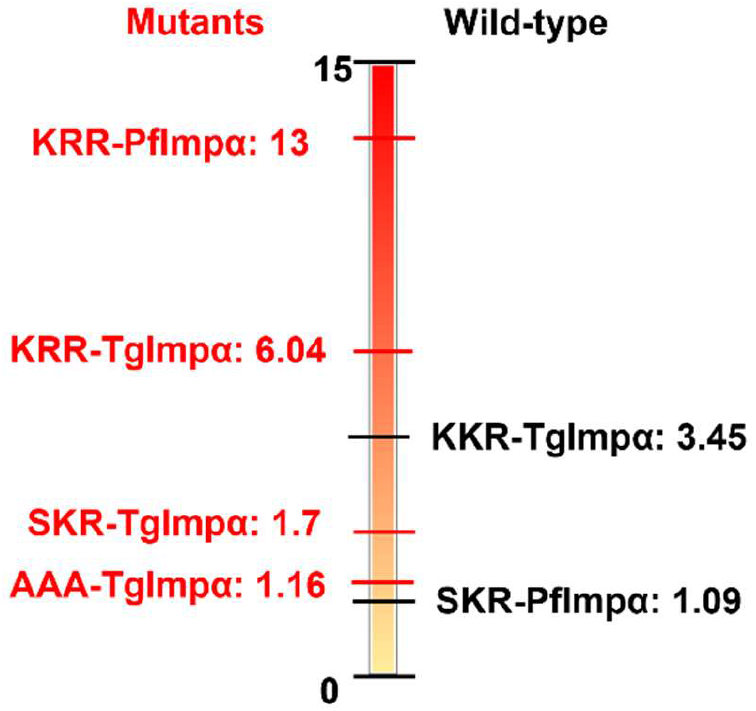
Comparison of wild-type and third basic cluster mutants of importin α from different organisms. Pf: *P. falciparum*, Tg: *T. gondii*. The fold-change values for binding the wild-type or mutant TgImpα proteins to PfTGS1-NLS are compared to ΔIBB-TgImpα and depicted in the schematic to allow a direct, visual comparison with wild-type and mutant proteins from *P. falciparum* (Dey & Patankar, 2018). Wild-type proteins are shown in black, while the third basic cluster mutant proteins are shown in red. The residues in the third basic cluster are mentioned with the name of the protein.

Interestingly, the KRR-TgImpα mutant showed a fold-change value of 6.04 compared to a value of ~2-3 for the wild-type protein. These results indicate that the strength of auto-inhibition of the wild-type TgImpα protein, having a KKR motif, can be increased by mutation of a single lysine residue to generate a KRR motif and suggest that TgImpα may exhibit weak auto-inhibition. Additionally, comparing the KRR-TgImpα fold-change value (6.04-fold) to the fold-change value of the same mutation in PfImpα (13-fold) indicates that factors other than the third basic amino acid cluster appear responsible for tuning the strength of auto-inhibition in TgImpα.

### *In silico* docking of the TgImpα IBB domain at the major NLS-binding site provides insights into the auto-inhibition strength

In an attempt to understand the phenotype of auto-inhibition seen for TgImpα, we carried out *in silico* structural analyses of the interactions between the IBB domain and the ARM repeats. Indeed, in a structural study of MmImpα2 (PDB ID: 1IAL), intramolecular interactions were able to explain the auto-inhibition caused by the third basic residue cluster (KRR) (Kobe, 1999).

The predicted structure models of full-length TgImpα and MmImpα2 were initially compared. Their helical ARM repeats and NLS-binding sites were superimposed (Figure 5A), indicating that the prediction had resulted in a structure that could be used for further analyses. Notably, TgImpα harbors a longer hinge than MmImpα2, and the TgIBB domain was predicted to have a helical structure (Figure 5A).

**Figure 5.**
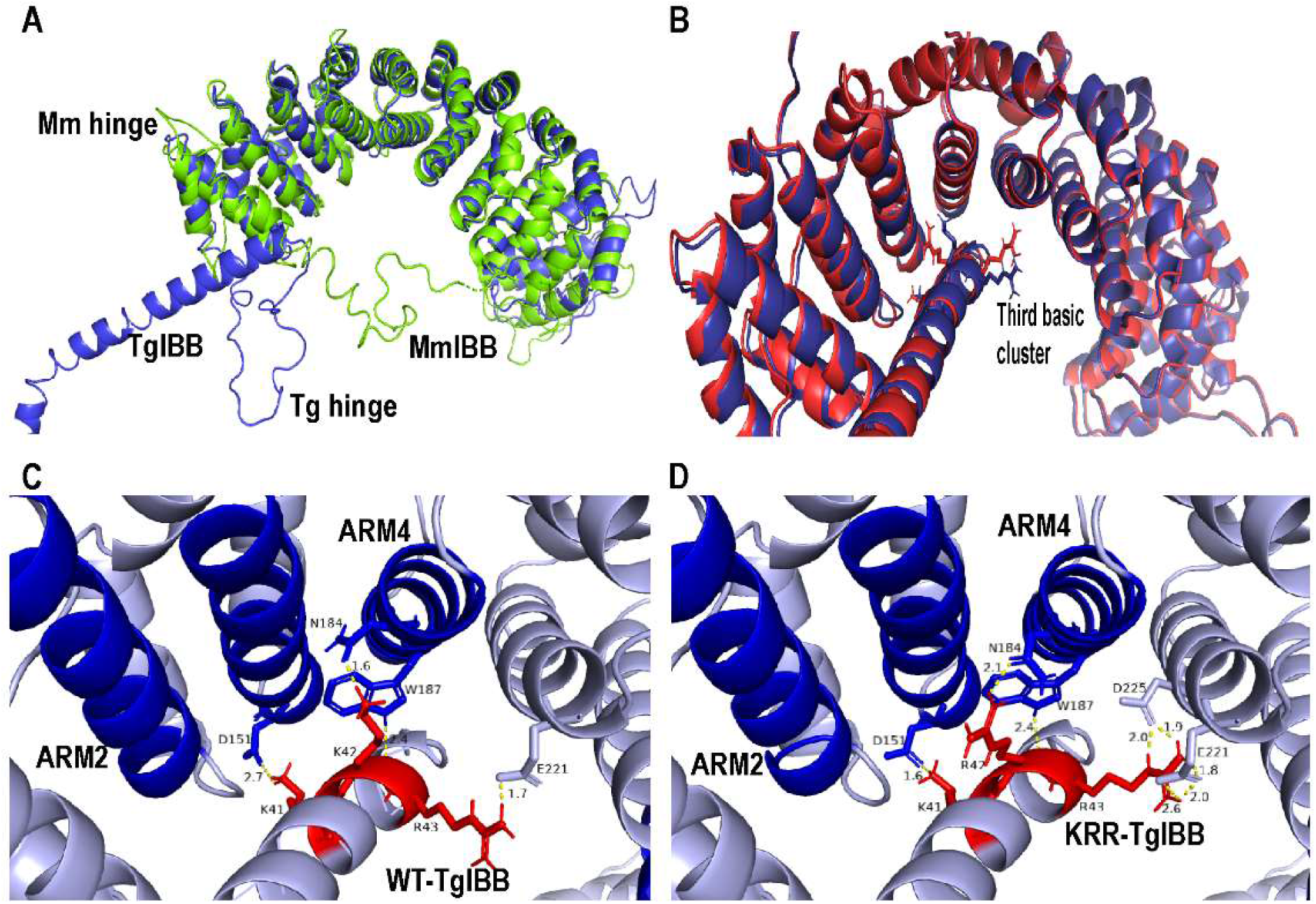
*Ab initio* models of the interaction at the IBB/ARM interface of TgImpα. **(A)** Predicted models of MmImpα2 (green) and TgImpα (blue) superimposed to show the similarity between the two predicted models at the ARM repeats but not at the IBB domain. A difference in the length of the hinge motif can be distinctly seen. **(B)** Superimposing predicted ΔIBB-TgImpα docked with predicted TgIBB (blue) and KRR-TgIBB (red), their third basic residue cluster and IBB are superimposed. Measurements of the polar bond distances at the IBB/ARM interface of the models when **(C)** TgIBB and **(D)** KRR-TgIBB are docked at ΔIBB-TgImpα, NLS-binding site (dark blue) and third basic cluster (dark red) are shown. Note that TgIBB denotes the WT-TgIBB protein without any mutations.

Next, residues 1-49 of the IBB domain of TgImpα (TgIBB) were modeled (estimated RMSD: 2.37 Ǻ). Also, they showed a helical structure (Figure 5B) that was predicted to have a helical content of 20.41% predicted by the Agadir program. Similar results were seen using the Robetta software and in the AlphaFold database. In contrast, the MmImpα2 IBB domain shows a lower helical content (11.87%), and the third basic cluster residues that form key contacts with the major NLS-binding site are in an unstructured motif (PDB ID:1IAL) (Kobe, 1999).

The prediction of a helical IBB domain and the longer hinge could potentially change the position and orientation of the third basic cluster, affecting the intramolecular interaction of TgIBB with the ARM repeats. In the docking of TgIBB and KRR-TgIBB estimated RMSD: 2.04 Ǻ) at the ARM repeats of ΔIBB-TgImpα (estimated RMSD: 5.21 Ǻ), differences in the orientation of the third basic residue cluster were observed (Figure 5B). Therefore, a detailed analysis of the interactions between the amino acid residues in the third basic residue cluster and the NLS-binding sites of the ARM repeats was performed.

The interaction between the third basic cluster, KRR, in the IBB domain of MmImpα2 and the ARM repeats is stabilized by hydrogen bonds with the NLS-binding site and the crystal’s water molecules (Kobe, 1999). These interacting residues and their polar bond distances are summarized in Table 1 to compare with the interactions observed in the predicted TgImpα models. In the docking analysis of the models, fewer interactions are observed between ΔIBB-TgImpα and TgIBB than those seen for MmImpα2 (Figure 5C). The summarised bond distances indicate that residues further away from the basic cluster also form interactions (Table 1). Compared to wild-type TgImpα, the KRR-TgImpα mutant demonstrates 1.7-fold stronger auto-inhibition, showing a stronger intramolecular interaction between the IBB domain and NLS-binding site. To validate this observation with the predicted model, we docked the ΔIBB-TgImpα and the KRR-TgIBB mutant structures (Figure 5D) and first observed the wild-type and mutant IBB domains were superimposing at the third basic cluster (Figure 5B). Despite this, changing single lysine to arginine (KKR to KRR) resulted in numerous new interactions of the IBB domain with the ARM repeats. For example, R43 in KRR-TgIBB forms polar interactions with the second and fourth ARM repeats but is not seen in the wild-type TgIBB protein. The R42 residue of the KRR mutant forms a polar bond with N184 of the ARM repeat. Residues flanking the third basic residue cluster of the IBB domain (N38 and D48) also showed new interactions with the ARM repeats in the KRR mutant (Table 1). These results also reinforce the data (Figure 4), suggesting that the wild-type TgImpα protein, having a KKR motif, shows weak auto-inhibition.

**Table 1.** Polar interactions formed at the interface between the IBB domain and the ARM repeats. Mm: *M. musculus*, Tg: *T. gondii*. The polar bond distances in Angstroms (Å), hy (hydrophobic) are in brackets with the residue number. The polar bond distances for MmImpα2 are from the PDB: 1IAL, and for TgImpα, wild-type and mutant IBB are from the docking analyses. Note that TgIBB denotes the WT-TgImpα IBB domain. Underlined are the third basic cluster residues.

| **MmImpα2 (PDB: 1IAL)** |  | **WT-TgIBB and ΔIBB-TgImpα** |  | **KRR-TgIBB and ΔIBB-TgImpα** |  |
| --- | --- | --- | --- | --- | --- |
| **IBB** | **ARM** | **IBB** | **ARM** | **IBB** | **ARM** |
| K42 |  | H34 | R61 (2.8), S64 (2.8) | H34 |  |
| K43 |  | R35 |  | R35 |  |
| D44 | W273 (hy), R315 (hy) | E36 |  | E36 |  |
| E45 | W273 (hy) | Q37 | S64 (2.5) | Q37 | S64 (2.2) |
| Q46 | S234 (2.8), D270 (2.5), Y277 (2.8), R238 (2.7) | N38 |  | N38 | W101 (2.1), N105 (2.1) |
| M47 | W231 (3.1) | L39 |  | L39 |  |
| L48 | N235 (3.2) | A40 |  | A40 |  |
| <u>K49</u> | G150 (3.0), T151 (3.3), T155 (2.8), D192 (2.7) | <u>K41</u> | D151 (2.7) | <u>K41</u> |  |
| <u>R50</u> | W184 (2.9), N188 (2.8, 2.9), N228 (2.9, 3.0) | <u>K42</u> | W187 (2.4) | <u>R42</u> | N184 (2.1), W187 (2.4) |
| <u>R51</u> | L104 (3.0), R106 (3.0, 3.0) | <u>R43</u> |  | <u>R43</u> | E221 (2.0, 2.6), D225 (2.0) |
| N52 | S105 (2.9), W142 (3.0), N146 (2.8, 2.9) | A44 |  | A44 |  |
| V53 |  | E45 | G150 (2.8) | E45 | N147 (2.1) |
| S54 |  | A46 |  | A46 |  |
| S55 |  | L47 |  | L47 |  |
| F56 |  | D48 | N191 (2.6) | D48 | R156 (2.0), R194 (2.3) |

### The hinge motif in importin α proteins of Apicomplexa is longer than that of other phyla

In the MSA of full-length importin α proteins from different phyla (Figure 3A), we observed different hinge motif lengths and plotted the hinge length in 73 importin α proteins from different organisms. Importin α proteins from most of the phyla showed a mean length of the hinge motif in the range of 15-20 amino acids. Interestingly, the hinge in the apicomplexan importin α proteins is almost twice as long as the hinge seen in other phyla, with a mean of >30 residues (Figure 6A). The length of the TgImpα hinge is 30 amino acids long, also observed in the predicted structure of full-length TgImpα (Figure 5A).

**Figure 6.**
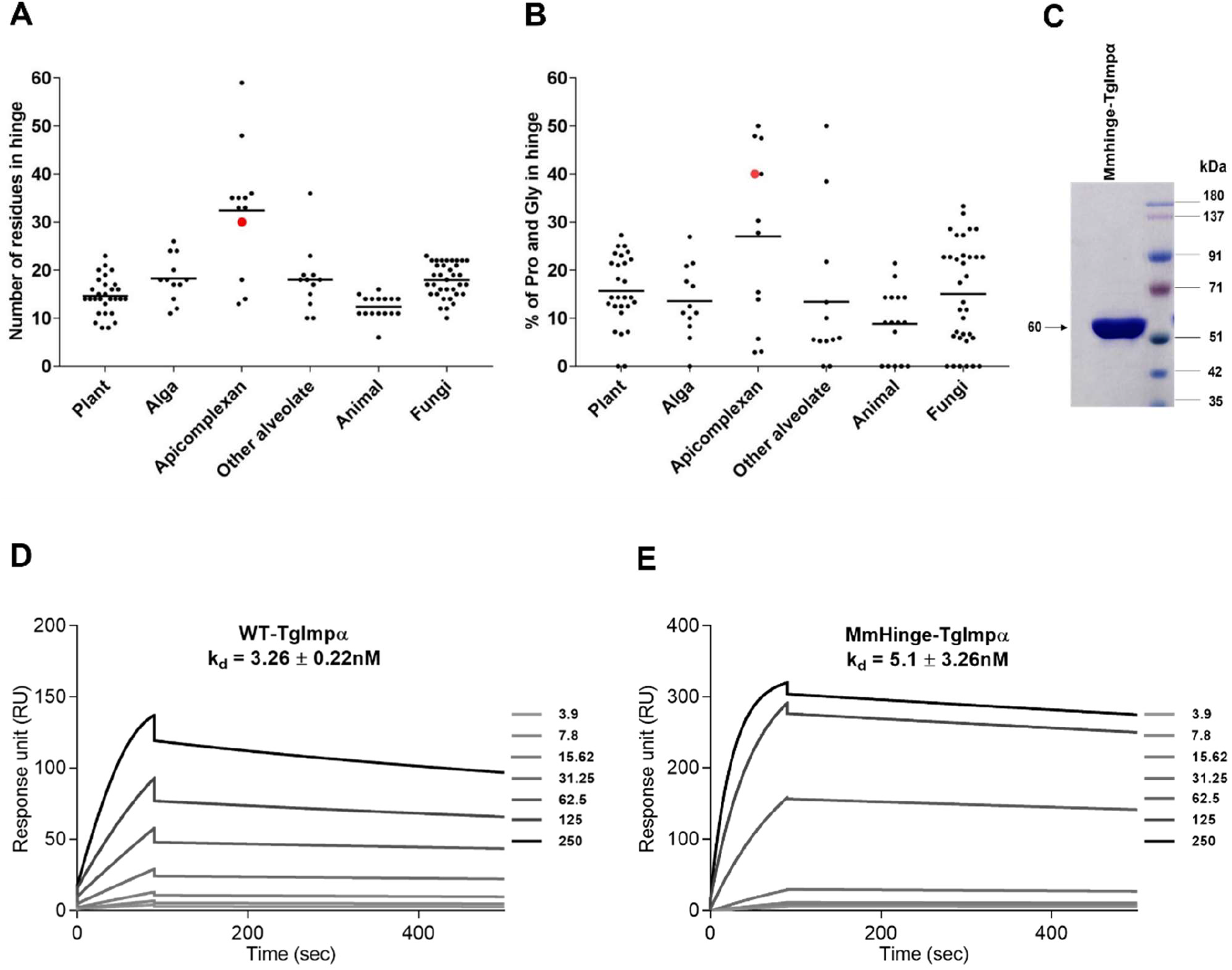
Hinge motif in importin α from different phyla varies in length and composition yet does not contribute to auto-inhibition of TgImpα. **(A)** Every dot is an importin α protein for the respective number of residues in the hinge motif with the mean value of the respective phyla. A significant difference is observed amongst groups, P < 0.0001 by ANOVA test. **(B)** Every dot is an importin α for the composition of proline and glycine residues in the hinge (values in percentage of the hinge length). A significant difference is observed amongst groups, P < 0.002 by ANOVA test. The dataset includes 73 proteins, the same as the phylogenetic analyses (see methods section). Highlighted is the hinge of TgImpα in red. **(C)** SDS-PAGE of purified MmHinge-TgImpα protein with an expected size of 60 kDa. SPR kinetic analysis of the interaction of **(D)** WT-TgImpα and **(E)** MmHinge-TgImpα with PfTGS1-NLS-GFP. All results represent the mean ± SD (n = 3) for Kd values measured. The concentration range mentioned is in nM.

A report has suggested that the hinge provides flexibility to the *H. sapiens* importin α3 protein (Sankhala et al., 2017). This leads to the speculation that in apicomplexans, a longer hinge might affect the orientation and positioning of the third basic cluster at the NLS-binding site interface. This could occur due to the flexibility of the hinge motif, a property that is influenced by the presence of amino acids disrupting the secondary structure, like proline and glycine. These amino acids are found at higher frequencies in the hinge motifs between two dynamic domains (Grummt, 1998; Veevers et al., 2020). Apicomplexan importin α proteins showed a mean of 30% proline and glycine in their hinge motifs, higher than the other phyla studied here (Figure 6B). Notably, in TgImpα, proline and glycine make up 40% of the hinge (Figure 6B, highlighted in red), suggesting higher flexibility, possibly affecting the positioning of the third basic residue cluster described in the previous section.

In order to directly test whether the hinge motif of TgImpα has any role in modulating the strength of auto-inhibition, we performed SPR assays with WT-TgImpα and a chimeric protein (MmHinge-TgImpα) containing a replacement of the TgImpα hinge (amino acids 49-78) with the MmImpα2 hinge (amino acids 57-70) (Figure 6C). Note that the hinge motif of MmImpα2 is smaller and contains only one proline and no glycine residues (Figure 3A). A Kd of 3.26 ± 0.22 nM was obtained for WT-TgImpα (Figure 6D) and 5.1 ± 3.26 nM for MmHinge-TgImpα (Figure 6E) with PfTGS1-NLS-GFP, while the negative control (GFP) showed no binding. Compared to WT-TgImpα, the MmHinge-TgImpα chimera does not demonstrate a significant increase in the Kd. These results indicate that the long, proline and glycine-rich hinge motif of TgImpα is not involved in auto-inhibition and may have alternate functions.

### Quantitative analysis of auto-inhibition of importin α proteins from the literature

In this report, the auto-inhibition strength is calculated as a fold-change value between the Kd of the wild-type protein and the protein lacking auto-inhibition. The fold-change value reflects the ability of the full-length importin α protein to bind to the NLS and higher values indicate that the protein shows strong auto-inhibition. In contrast, a fold-change value of 1 indicates that the full-length importin α protein binds with similar affinity to the NLS as the ΔIBB protein or an assay including importin β, suggesting a lack of auto-inhibition. We compiled published data to compare fold-change values of different importin α proteins and assess whether the strength of auto-inhibition can be inferred from these data (Table 2).

**Table 2.**
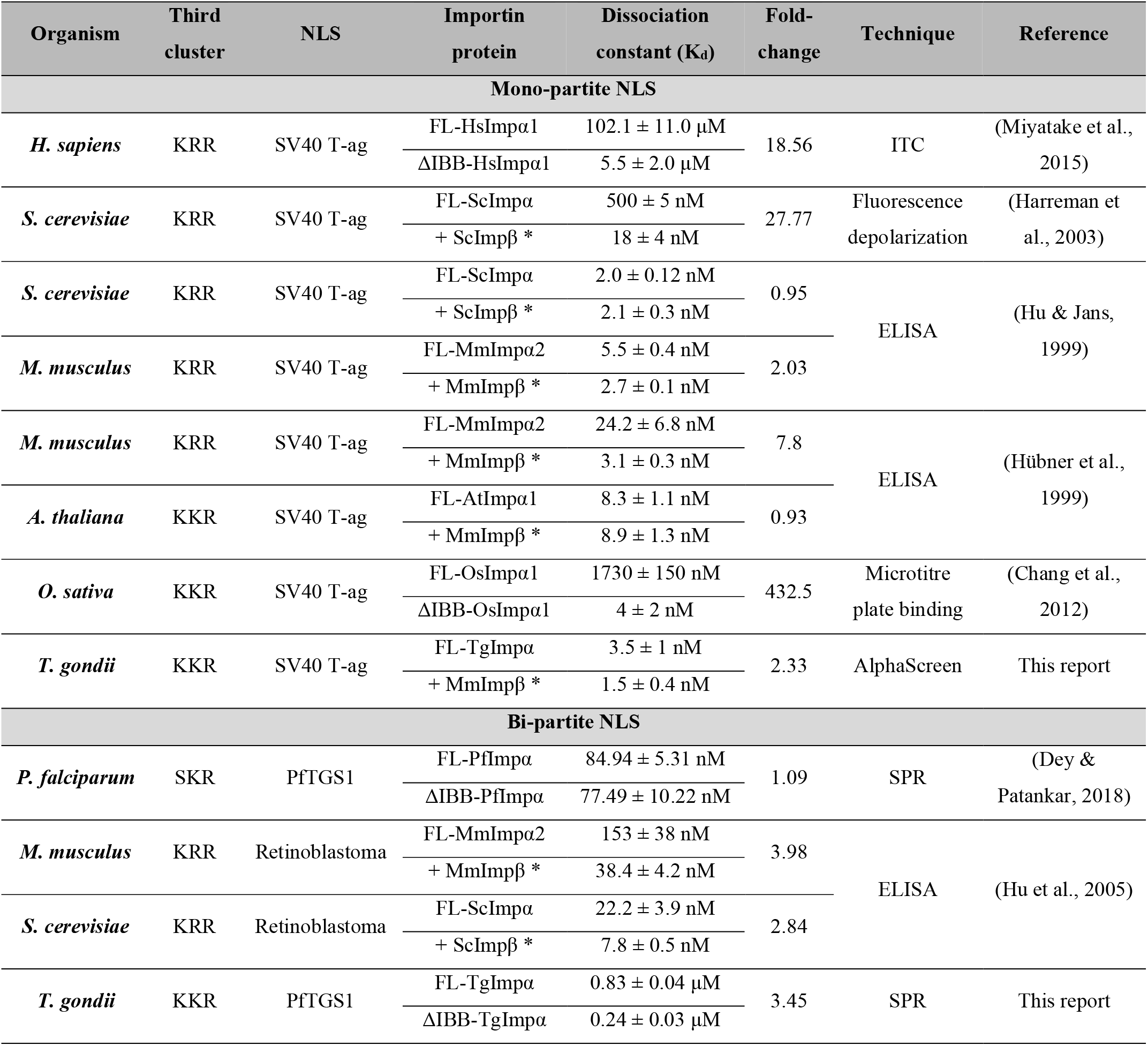
Auto-inhibition of importin α proteins from different organisms. The sequence of the third basic cluster in the IBB domain, the NLS used in binding assays and the dissociation constants of the importin α from different organisms are shown. The fold-change between the Kd of the full-length (FL) importin α and the protein that lacks auto-inhibition, either due to IBB domain deletion (ΔIBB) or due to the addition of importin β (shown with an asterisk), is calculated and serves as a measure of the strength of auto-inhibition. The experimental techniques and types of NLSs are mentioned. ITC: Isothermal calorimetry; ELISA: Enzyme-linked immunosorbent assay; SPR: Surface plasmon resonance.

Analysis of quantitative data from the literature (Table 2) shows that the Kd values can change depending on the assay and the NLS. Clearly, to compare Kd values and strength of auto-inhibition between different importin α proteins, they must be assayed in the same experiment. However, these data also indicate that the fold-change values of importin α proteins that show auto-inhibition is consistently greater than 1, regardless of the assay and the NLS being mono-partite or bi-partite, except for one study where the fold-change value for *S. cerevisiae* importin α is unusually low (Hu & Jans, 1999). The results presented in this report, when set against the compiled data from Table 2, reinforce the conclusion that *T. gondii* importin α shows auto-inhibition, unlike *P. falciparum* importin α, which shows a fold-change value of 1.09 and, therefore, a lack of auto-inhibition. *A. thaliana* importin α1 has a fold-change value of 0.93, also showing a lack of auto-inhibition (Table 2). Interestingly, this protein has a KKR motif in the third basic residue cluster.

### Apicomplexan importin α proteins are not evolutionarily related to those from plants

Multiple lines of evidence show the inheritance of an extant non-photosynthetic plastid, the apicoplast, in apicomplexans from a common red algal endosymbiont (Janouskovec et al., 2010). Indeed, a minimal phylogenetic tree consisting of five importin α proteins, PfImpα, TgImpα, AtImpα, ScImpα and HsImpα, showed that apicomplexan importin α proteins are more similar to those found in plants (Bhatti & Sullivan, 2005). We asked whether an extensive phylogenetic analysis would also show that apicomplexan importin α proteins had an evolutionary history that could be traced to the endosymbiont. If this were the case, the phylogenetic analysis would place the apicomplexan importin α proteins closer to algae and plants.

Apicomplexans, dinoflagellates, colpodellids, ciliates and perkinsozoans are members of the Alveolata group (Van de Peer & De Wachter, 1997). The apicoplast in apicomplexans shares a common origin with the plastids of dinoflagellates and colpodellids, the closest algal relatives (Fast et al., 2001; Janouskovec et al., 2010, 2019). However, during our analysis, dinoflagellate importin α amino acid sequences were highly divergent when aligned through the MUSCLE program and were removed. Among the 5 colpodellid importin α proteins, only two *Vitrella brassicaformis* importin α proteins aligned in MUSCLE and branch with the apicomplexan clade, indicated by a branch of Apicomplexa and Colpodellida in the tree (Figure 7, highlighted in blue).

**Figure 7.**
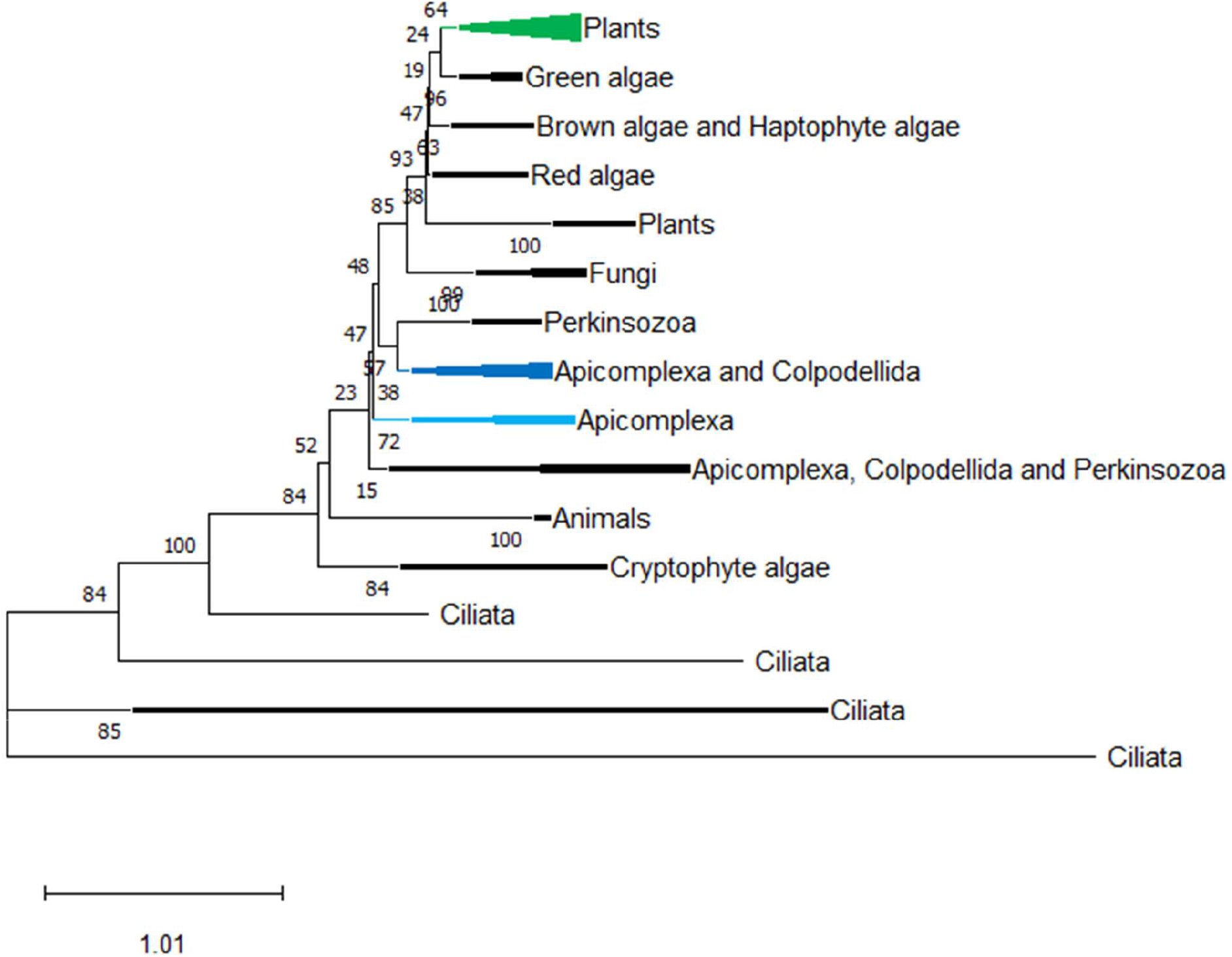
Importin α of apicomplexans *P. falciparum* and *T. gondii* and plant *A. thaliana* shows no evolutionary relatedness. Maximum likelihood phylogeny of the importin α proteins. The highlighted branches are AtImpα1 (green), TgImpα (blue) and PfImpα (cyan). The other sequences are from *Theileria, Babesia, Cryptosporidium, Neospora, Besnoitia, Cystoisospora, Cyclospora, Gregarina, Hepatocystis* and *Eimeria* (apicomplexan); *Chlamydomonas, Volvox*, *Chlorella* and *Auxenochlorella* (green alga); *Ectocarpus* (brown alga); *Emiliania* (haptophyte alga); *Galdieria, Porphyra* and *Gracilariopsis* (red alga); *Saccharomyces, Schizosaccharomyces* and *Neurospora* (fungi); *Perkinsus* (perkinsozoan); *Vitrella* (colpodellid); *Guillardia, Hemiselmis* and *Chroomonas* (cryptophyte alga); *Paramecium* and *Tetrahymena* (ciliate); *Homo* and *Mus* (animal). The apicomplexans and ciliates importin α branched into multiple clades. The dataset used for the phylogenetic analysis contained 73 importin α proteins. The branching pattern was collapsed to show representative clades and not individual organisms.

The Apicomplexa group branched with the fungi and plant group and not animals. In our analysis, apicomplexan importin α seems to be closely related to fungi and red algae (through the ancestral endosymbiont) and distantly to plants, a group containing *A. thaliana* (Figure 7, highlighted in green). Therefore, this extensive analysis with a diverse dataset does not indicate a close evolutionary relationship between apicomplexans (highlighted in blue and cyan) and plants (highlighted in green) (Figure 7). Despite PfImpα and AtImpα1 showing a lack of auto-inhibition, each protein may have acquired phenotype independently.

## DISCUSSION

### *T. gondii* importin α shows auto-inhibition that may be weak compared to other organisms

In contrast to *P. falciparum, T. gondii* importin α demonstrates auto-inhibition; this was confirmed with different techniques, NLSs and approaches. We propose that this auto-inhibition may be weak compared to other organisms. When the strength of auto-inhibition of TgImpα is quantified by calculating the fold-change value of binding affinities between the full-length protein and the protein lacking auto-inhibition, this value is ~2-3 fold. Although it is tempting to suggest that this number is lower than the fold-change values seen for other importin α proteins, we show in this report that rather than an intrinsic property of the protein, different NLSs and assays can give different fold-change values.

What, then, is the evidence for weak auto-inhibition? The strength of auto-inhibition of wild-type *T. gondii* importin α can be increased almost 2-fold by a single amino acid mutation in the third basic cluster of the IBB domain. Although both motifs have basic amino acids, a KKR motif appears to result in weaker auto-inhibition than a KRR motif. Using *in silico* structural analyses of wild-type and the KRR mutant of TgImpα, we show that the interactions between key amino acid residues of the NLS-binding sites and the KKR motif are strengthened in the KRR mutant. The hydrogen bonding of certain residues is energetically favorable over others at the NLS-binding pocket resulting in strong interactions (Conti et al., 1998). Our *in silico* analysis suggests that the NLS-binding pocket in TgImpα could favor arginine over a lysine residue in this motif. Identifying a mutant that increases the auto-inhibition strength provides a tool for testing the role of auto-inhibition *in vivo*.

The strength of auto-inhibition could also depend on other features in the protein as observed by *in silico* prediction analysis, the structure of the IBB domain of *T. gondii* importin α hints that this domain could be helical. The available forms of the IBB domain in complex with different importin proteins show helical structures in some cases and an unstructured domain in others, indicating structural polymorphism of the IBB domain (Cingolani et al., 1999, Matsuura & Stewart, 2004, Jibiki et al., 2021). Nevertheless, our structure prediction analyses indicate that in direct comparison to MmImpα2, there appears to be a higher α-helical content in the IBB domain of TgImpα. This structured IBB domain may play a role in the difference in the auto-inhibition strength between wild-type TgImpα and the KRR mutant.

Another interesting observation from the MSA was that the importin α proteins of apicomplexans harbor a longer hinge, almost twice the length seen in other taxa and a high frequency of prolines and glycines that disrupt helical structures. Importin α is a helix of helices with a strongly conserved rigid structure (Kobe, 1999; Pumroy et al., 2015; Sankhala et al., 2017). In such a rigid structure, the intramolecular interactions between the IBB and ARM domains are facilitated by the unstructured hinge motif that folds the IBB upon itself. However, our results show that the long hinge region of TgImpα does not play a role in the strength of auto-inhibition and could have other functions, such as interactions with regulatory proteins. Nevertheless, detailed structural studies of the apicomplexan importin α proteins will shed light on the helical content of the TgImpα IBB domain and the length and composition of the hinge motif as a basis for the strength of auto-inhibition.

### Phylogenetic analysis of importin α proteins reveals relationships between apicomplexans and single-celled eukaryotes

It was noteworthy that a lack of auto-inhibition is seen in the importin α proteins of *P. falciparum* and *A. thaliana* (Table 2) (Dey & Patankar, 2018; Hübner et al., 1999). These organisms share an evolutionary history as the extant non-photosynthetic plastid in apicomplexans may have been inherited from a common ancestor (Janouskovec et al., 2010). Previous work suggests that the importin α proteins from plants and apicomplexans are close relatives (Bhatti & Sullivan, 2005). We performed a detailed study by including other importin α homologs in our dataset. Interestingly, the phylogenetic tree with all phyla suggests a divergence between the plants and apicomplexans and a closer relation with red algae and fungi. We speculate that the same phenotype might have been acquired independently to suit each organism’s requirements for nuclear transport.

The evolutionary analysis of importin α proteins gave some novel insights. The branching of these proteins in Colpodellida and Perkinsozoa with Apicomplexa suggests a strong similarity. However, the ciliates branch at the bottom of the tree and diverges from all other alveolates. Our analysis of the three basic clusters in the IBB domain shows a lack of basic residue clusters in the IBB domain in ciliates (data not shown). The ciliates are known to have two nuclei (macronuclei and micronuclei) (Hausmann & Bradbury, 1996), and differential permeability regulates the nuclear import because of unique nucleoporins in their NPCs and importin α proteins specific to a single nucleus (Malone et al., 2008). These unique features may underlie the high variability in their IBB domain.

### Conclusions

In summary, we demonstrate that the auto-inhibition strength of *Toxoplasma gondii* importin α is conferred mainly by the third basic residue cluster in the IBB domain. We generate and characterize single amino acid mutants in the third basic residue cluster, resulting in a lack of or increased auto-inhibition strength. These mutants can be tested further using both structural and *in vivo* approaches to understand the role of auto-inhibition of *T. gondii* importin α in the biology of this important human pathogenic parasite.

## ACKNOWLEDGEMENTS

SP acknowledges the Department of Science and Technology, Science and Engineering Research Board, India, for funding (CRG/2018/000129). PhD fellowships were given to MB, VD and SW by the Human Resource Development Group-Council of Scientific and Industrial Research, Department of Biotechnology-The World Academy of Sciences and Indian Institute of Technology Bombay-Monash Research Academy. We acknowledge the SPR central facility funded by Industrial Research and Consultancy Centre at IIT Bombay and FP7 WeNMR, H2020 West-Life, EOSC-hub and EGI-ACE European e-Infrastructure projects for the docking studies. We thank Kiran Kondabagil, IIT Bombay, and David Jans, Monash University, for their valuable feedback on the manuscript.

## AUTHORS’ CONTRIBUTION

VD, SW, MB and SP conceived and designed the experiments. VD, SW and MB performed the *in vitro* experiments. MB and SP conceived and MB performed the *in silico* analysis. MB prepared the figures. MB and SP prepared the manuscript. MB, SP, VD and SW reviewed drafts of the manuscripts.

